# Healthy ageing influences how the shape of alpha and beta oscillations change during reaction time tasks

**DOI:** 10.1101/2023.10.16.562636

**Authors:** George M Opie, James M Hughes, Rohan Puri

**Author notes:** Correspondence: George M. Opie.

## Abstract

Age-related changes to the power and frequency of the brains oscillatory activity have been reported by an extensive literature. In contrast, the influence of advancing age on the *shape* of oscillation waveforms, a characteristic with increasingly recognised physiological and functional relevance, has not been previously investigated. To address this, we examined the shape of alpha and beta band oscillations from electroencephalography (EEG) data recorded during performance of simple and go/no-go reaction time tasks in 33 young (23.3 ± 2.9 years, 27 females) and 27 older (60.0 ± 5.2 years, 23 females) adults. The shape of individual cycles was characterised using instantaneous frequency, and then decomposed into waveform motifs using principal component analysis. This analysis identified four principal components (one from the alpha band, 3 from the beta band) that were uniquely influenced by the different motor tasks and/or age. These each described different dimensions of shape and tended to be modulated during the reaction phase of each task. However, the way in which each facet of shape varied during the task was unrelated to motor performance, indexed via reaction time, in either group or band. Our results suggest that although oscillation shape is task-dependent, the nature of this effect is altered by advancing age. While these outcomes demonstrate the utility of this approach for understanding the neurophysiological effects of ageing, future work that more clearly links these outcomes with function will be critical.

## Introduction

While the ageing process is associated with many changes to functional capacity, deficits within the motor system can have some of the most significant impact on independence and quality of life. Although the factors that drive these deficits remain poorly understood, there is good evidence that changes in brain dynamics are an important element. In particular, numerous studies have examined how the brain’s oscillatory activity – which is seen as rhythmic fluctuations in its electrical potential/magnetic field strength – is altered by age. This literature suggests substantial changes associated with ageing, including reductions in power and peak frequency within the alpha band (∼8-13 Hz; Chiang *et al*. 2011, Barry and De Blasio 2017, Scally *et al*. 2018, Sghirripa *et al*. 2021, Merkin *et al*. 2023, Tröndle *et al*. 2023), in addition to increased power (Rossiter *et al*. 2014, Heinrichs-Graham and Wilson 2016, Barry and De Blasio 2017, Heinrichs-Graham *et al*. 2018, Rempe *et al*. 2022) and frequency (Zhong and Chen 2022) in the beta band (∼14-30 Hz). Importantly, these changes have been associated with deficits in motor function, including reduced skill learning (Rueda-Delgado *et al*. 2019), reaction time (Van Hoornweder *et al*. 2022b) and accuracy (Van Hoornweder *et al*. 2022a). Furthermore, altered oscillatory activity has also been associated with motor pathologies common in older adults, such as Parkinson’s disease (Little *et al*. 2012, De Hemptinne *et al*. 2013, De Hemptinne *et al*. 2015, Cole *et al*. 2017). Age-related changes in oscillatory activity therefore appear to be a functionally relevant element of ageing, and may have potential as biomarkers of age-related degradation, or accumulating pathology.

Although effects of age on oscillatory activity are supported by numerous studies, outcomes have also been variable. For example, previous work has failed to replicate the often reported age-related reduction in alpha frequency (Polich 1997, Gaál *et al*. 2010, Caplan *et al*. 2015, Zhong and Chen 2022), whereas effects of age on both frequency and amplitude may be confounded by age-related changes in non-oscillatory activity also present in EEG recordings (Donoghue *et al*. 2020, Merkin et al. 2023, Tröndle et al. 2023). This variability limits the utility of oscillatory metrics for understanding the ageing process and challenges their clinical application. One factor that may contribute to this variability is the way in which oscillatory activity has been assessed. The conventional approach to quantifying neuronal oscillations generally involves characterising their frequency or amplitude via Fourier-based methods applied to time series data that generally involve several minutes of recordings. However, this approach overlooks important features of oscillations that are only apparent in the time domain. In particular, the developing literature shows that examination of waveform *shape* can provide information that is physiologically and functionally relevant. For example, recent work suggests that a given oscillatory cycle can be characterised relative to a range of waveform motifs, with cycle-by-cycle variations in the relative contribution of different motifs driving variability in shape (Quinn *et al*. 2021b, Szul *et al*. 2022, Rayson *et al*. 2023) and having different relationships with movement (Szul et al. 2022, Rayson et al. 2023) and function (Quinn et al. 2021b). Furthermore, research in Parkinson’s disease supports the clinical utility of examining waveform shape. Specifically, the shape of beta oscillations is significantly altered in patients off medication (Cole et al. 2017, Jackson *et al*. 2019), but this is corrected by medication (Jackson et al. 2019) or deep brain stimulation (Cole et al. 2017).

While the developing literature demonstrates the utility of examining the shape of oscillatory activity at the level of individual cycles, it remains to be investigated if this approach is sensitive to the neurophysiological and functional changes associated with advancing age. Within the current study, we aimed to address this limitation. The shape of oscillatory activity recorded at rest, or while performing a simple or go/no-go reaction time task, was quantified and compared between young and older adults. This was achieved by using a recently established methodology (Quinn et al. 2021b), within which empirical mode decomposition (EMD) facilitates extraction of multiple narrowband waveforms for individual cycles of the target oscillation. Subsequent application of principal component analysis (PCA) then identifies dominant waveform motifs. Given the well-established effects of ageing on conventional measures of oscillatory activity within the alpha and beta bands (see above), these served as the bands of interest within the current study.

## Methods & Methods

### Dataset

The electroencephalography (EEG) recordings analysed in the current study were obtained from a recently described open access dataset (Ribeiro and Castelo-Branco 2019). This study recruited 36 young (mean age ± SD: 23.1 ± 2.8 years; 29 females) and 39 older (mean age ± SD: 60.4 ± 5.2 years; 31 females) adults to participate in a single session, within which EEG was recorded in different conditions (see below). A number of participants within this dataset presented signal noise that would have required low-pass filtering to remove. As this type of processing can be expected to influence the shape of the EEG recording (de Cheveigné and Nelken 2019), and given our primary interest in quantifying waveform shape, we therefore decided to exclude these participants from the analyses. Consequently, the current study examined data from a subset of 60 participants, including 33 young (mean age ± SD: 23.3 ± 2.9 years; 27 females) and 27 older (mean age ± SD: 60.0 ± 5.2 years; 23 females) adults. All experimentation was performed in accordance with the Declaration of Helsinki, participants provided written, informed consent prior to inclusion, and the protocol was approved by the Ethics Committee of the Faculty of Medicine at the University of Coimbra.

### Experimental Task

Participants completed a cued reaction time task, which required a button press with the right index finger (or no response, depending on the condition) in response to auditory stimuli. A passive listening condition involving 30 trials was first completed, wherein participants were exposed to the auditory tones to be used in the active conditions but were not required to respond (hereafter referred to as ‘rest’). Following this, simple (SRT) and go/no-go (GNG) reaction time tasks were completed, the order of which was counterbalanced between participants. During the SRT, a cue tone (indicating the start of the trial) was followed at a variable interval by a ‘go’ tone for a total of 100 trials. During the GNG, a cue tone was followed at a variable interval by either a ‘go’ tone, requiring a button press response (80 trials), or a ‘no-go’ tone, requiring participants to withhold a response (20 trials). For both SRT and GNG, 20 catch trials were also included, in which the cue tone was not followed by any further tone. Between trials, participants fixated on a cross displayed on a screen in front of them. Slow trials were defined as a reaction time exceeding 700 ms, feedback on which was provided to participants by a different tone. The inter-trial interval ranged from 6.7 s to 19.6 s, with a median value of 7.6 s.

### Electroencephalography (EEG) acquisition and pre-processing

EEG was recorded with a Neuroscan system via 64 electrodes in standard 10-20 locations. The signal was referenced to a location between CPz and Cz, the ground was located between FPz and Fz, and data were digitized at a rate of 500 Hz. Pre-processing used custom scripts on the Matlab platform (R2021b, Mathworks, USA) with EEGLAB (v2022.1)(Delorme and Makeig 2004) and TESA (v 1.1.1)(Rogasch *et al*. 2017) toolboxes. Slow drifts in the signals were first removed by high-pass filtering above 1 Hz using the *pop_eegfiltnew* function. Line noise and its first harmonic (i.e., 50 & 100 Hz) was then attenuated using the EEGLAB CleanLine plugin (Mullen 2012), which uses a multi-tapering approach to remove line noise while minimising signal distortion. Data were then epoched from 500 ms before to 6000 ms after the cue tone. Channels and epochs demonstrating persistent, large amplitude muscle activity or noise were then removed. Following this, independent component analysis (ICA) was run using the FastICA algorithm (Hyvärinen and Oja 2000) and components associated with blinks, muscle activity, eye movement, and electrode noise were identified and removed based on visual inspection of component time course and topography. Missing channels were then replaced using spherical interpolation.

### Waveform analysis

All subsequent analysis of EEG data focussed on the C3 electrode, given its assumed location over the left sensorimotor cortex (Lefaucheur *et al*. 2017) activated during performance of reaction time tasks involving the right index finger. Furthermore, to facilitate comparisons with the SRT task, only trials from the ‘go’ condition of the GNG task were included in the analysis. Analysis of waveform shape of individual oscillatory cycles was performed according to the pipeline developed recently by Quinn et al. (2021b). This involved: (1) application of empirical mode decomposition (EMD) to decompose the recorded broadband signal into discrete narrowband oscillatory modes; (2) identification of individual cycles within oscillations of interest, and phase alignment to allow comparisons of shape between cycles with varying temporal dynamics and (3) application of principal component analysis (PCA) to identify consistent variations in cycle shape as waveform motifs. All analyses were performed in Python 3.10, using v0.4.0 of the EMD package (Quinn *et al*. 2021a) and v1.4.0 of the SAILS package (Quinn and Hymers 2020).

#### Empirical mode decomposition

EMD uses an iterative sifting process to decompose a broadband signal into narrowband intrinsic mode functions (IMFs), whereby higher frequency components of the signal are progressively extracted and subtracted from the signal. Briefly, maxima and minima of the broadband signal are identified, and upper and lower envelopes of the signal are developed by connecting and interpolating the maxima and minima, respectively. The mean of the overall envelope is then calculated and subtracted from the signal. This process is repeated on the resulting waveform until the criteria defining an IMF are met. These are that the number of zero crossings equals the number of extrema (differing by no more than 1) and the mean of the signal envelope equals zero (Huang *et al*. 1998). The first IMF is then subtracted from the original signal and the process is repeated to find the next IMF. Unlike conventional approaches to generating a narrowband signal that assume a sinusoidal waveform, this process conserves the native shape of the target oscillation (Quinn et al. 2021b).

Within the current study, the EMD algorithm was applied using previously established options (Quinn et al. 2021b) and a maximum of 6 IMFs were generated. We applied the masked version of EMD, where a sinusoidal masking frequency is added to the waveform prior to sifting (Deering and Kaiser 2005). This reduces the impact of mode mixing, which refers to situations in which different frequency components are mixed into a single IMF due to noise or intermittent oscillations in the signal (Huang *et al*. 1999). Mask frequencies of 120, 64, 32, 11, 7 and 2 Hz were applied, and validated by examination of instantaneous frequency profiles for each IMF (see supplementary figure S1); these showed clean separation between IMFs. Modes corresponding to conventional alpha and beta bands were found in the third (beta) and fourth (alpha) IMFs for all participants.

To demonstrate the reliability of the IMFs prior to examination of single cycles, age-related changes in alpha amplitude and peak frequency were compared between estimates derived from the broadband data and the alpha IMF. To increase sensitivity to effects of age on alpha activity, this analysis utilised data derived from the Oz electrode. Broadband and IMF data from individual epochs in the passive listening condition were concatenated to form individual time series, which were then decomposed using Welch’s method (8s window length, 50% overlap between windows). Power and frequency values associated with the alpha band (8-13Hz) within each group were then compared between techniques using independent samples t-tests, and the correlation between them was tested with Spearman’s rho.

#### Cycle detection and phase alignment

Following EMD, individual cycles in the alpha and beta modes were identified based on the instantaneous amplitude and phase of the signal, derived using the normalised Hilbert transform (Quinn et al. 2021b). Cycles were only included in further analyses if their amplitude exceeded the 50^th^ percentile for the mode (Fabus *et al*. 2021, Echeverria-Altuna *et al*. 2022), their phase characteristics met previously defined criteria for a reliable cycle, and they possessed unique control points (i.e., ascending/descending zero crossings, peak and trough)(Quinn et al. 2021b). The instantaneous frequency (IF) was then calculated for each cycle by taking the first derivative of instantaneous phase with respect to time (Huang *et al*. 2009). Waveform shape was then inferred from the IF of the cycle, wherein within-cycle fluctuations in IF indicate deviations in shape. However, it is not possible to directly compare IF profiles between different oscillations, as it is not clear if comparisons are occurring between consistent features of each cycle (e.g., comparing values from the peak of each cycle)(Quinn et al. 2021b). Identified cycles were therefore aligned to a common phase space, producing phase-aligned IF profiles that were directly comparable between oscillation cycles (Quinn et al. 2021b). To quantify the extent to which individual cycles deviated from a sinusoidal shape, phase-aligned IF profiles were projected onto the complex plane and a mean vector was calculated, with non-zero values indicating a non-sinusoidal shape. Real and imaginary values derived from individual cycle mean vectors (reflecting ascending-descending and peak-trough asymmetry, respectively) were compared to 0 using single sample t-tests (Quinn et al. 2021b).

#### Principal component analysis

The final step of the waveform analysis was to characterise the variability in waveform shape across cycles by submitting the phase-aligned IF profiles to PCA. The principal components (PCs) derived from this analysis can be viewed as waveform motifs describing major sources of variance relative to the mean instantaneous frequency profile. Furthermore, the scores describe the extent to which each PC contributes to a given cycle. The first 5 PCs within each frequency mode explained > 97% of variance in IF and were used for further analysis.

### Statistical analyses

Given recent evidence for temporal variance in waveform shape during a motor task (Szul et al. 2022, Rayson et al. 2023), a time factor was included for all analyses of cycle features. This categorised cycles as occurring either before (pre-go) or after (post-go) the go stimulus, or after the button press (post-react). Within each band, effects of task condition (rest, SRT*_pre-go_*, SRT*_post-go_*, SRT*_post-react_*, GNG*_pre-go_*, GNG*_post-go_*, GNG*_post-react_*) and group (young, older) on mean within-cycle phase aligned IF and PC score were assessed using Bayesian generalised linear mixed models (GLMMs), resulting in 12 models being run (i.e., one model for phase aligned IF and one model for each of the first 5 PCs, within both alpha and beta bands). Within each model, a student’s *t* distribution and an identity link function were applied. The maximal random effects structure allowed by the data was used (i.e., by-participant random intercepts and slopes)(Barr 2013). Potential relationships between PC scores and performance (indexed by reaction time [RT]) were then investigated. For each trial, PC scores within each time point (i.e., pre-go, post-go and post-react) were summarised using the coefficient of variation and median. These were then correlated with RTs using robust Bayesian correlations (https://baezortega.github.io/2018/05/28/robust-correlation/).

All posterior distributions were estimated with the No-U-Turn-Sampler (NUTS) extension of Hamiltonian MCMC, implemented within the BRMS package (Bürkner 2017). Each model was run using 4 independent chains with 1000 warm up and 3000 post-warm up samples (total of 12,000 post-warm up samples) and default flat priors. Chain convergence was assessed by ensuring Rhat values were < 1.1, in addition to visual inspection of post-warm up samples (Gelman and Rubin 1992). Posterior predictive checks were conducted to ensure simulated data matched observed data (Gabry *et al*. 2019). After model fitting, the *emmeans* package (Lenth 2023) was used to generate custom contrasts; these included within- and between-subject effects for all dependent variables, in addition to interaction contrasts where relevant. Within these comparisons, effect *existence* was described using the probability of direction (*pd*), which reports the proportion of the posterior with the same sign as the median and ranges from 50% to 100% (Makowski *et al*. 2019). Effect *significance* was assessed by examining how far the posterior distribution for each contrast deviated from a region including zero (i.e., no practical difference). To achieve this, a region of practical equivalence (ROPE) was first defined; this refers to a range of values centred around zero that would be considered as practically equivalent to no difference for that contrast. Within the current study, this was set as ± 5% of the standard deviation (SD)(Kruschke 2018). The 89% highest density interval (HDI; i.e., the range which contains 89% of the posterior distribution) was then identified, and the extent to which it overlapped the ROPE was used to make a decision regarding the null hypothesis. The null hypothesis of no difference was accepted if the 89% HDI fell completely inside the ROPE or rejected if it fell completely outside the ROPE. In contrast, no decision was made if the 89% HDI partially overlapped the ROPE. (Kruschke 2018, Puri *et al*. 2023).

## Results

As an initial step, measures of oscillatory activity provided by EMD were validated against those provided by conventional approaches to spectral analysis. This was achieved by applying Welch’s method to both the alpha mode and the broadband data, and comparing estimates of alpha frequency and amplitude. For both techniques, while there was no significant difference in peak frequency between groups (broadband: *t*_58_ = 1.0, *p* = 0.3; alpha IMF: *t*_58_ = 0.5, *P* = 0.6; Fig 1A2 & 1B2), power was significantly reduced in older adults (broadband: *t*_58_ = 2.8, *p* = 0.008; alpha IMF: *t*_58_ = 2.90, *P* = 0.005; Fig 1A3 & 1B3). Furthermore, estimates of both frequency (young: rho = 0.98, *p* < 0.0001; older: rho = 0.77, *p* < 0.0001) and power (young: rho = 0.99, *P* < 0.0001; older: rho = 0.98, *p* < 0.0001) were highly correlated between techniques (Fig 1C & 1D). These results support the reliability of the IMFs extracted by EMD.

**Figure 1.**
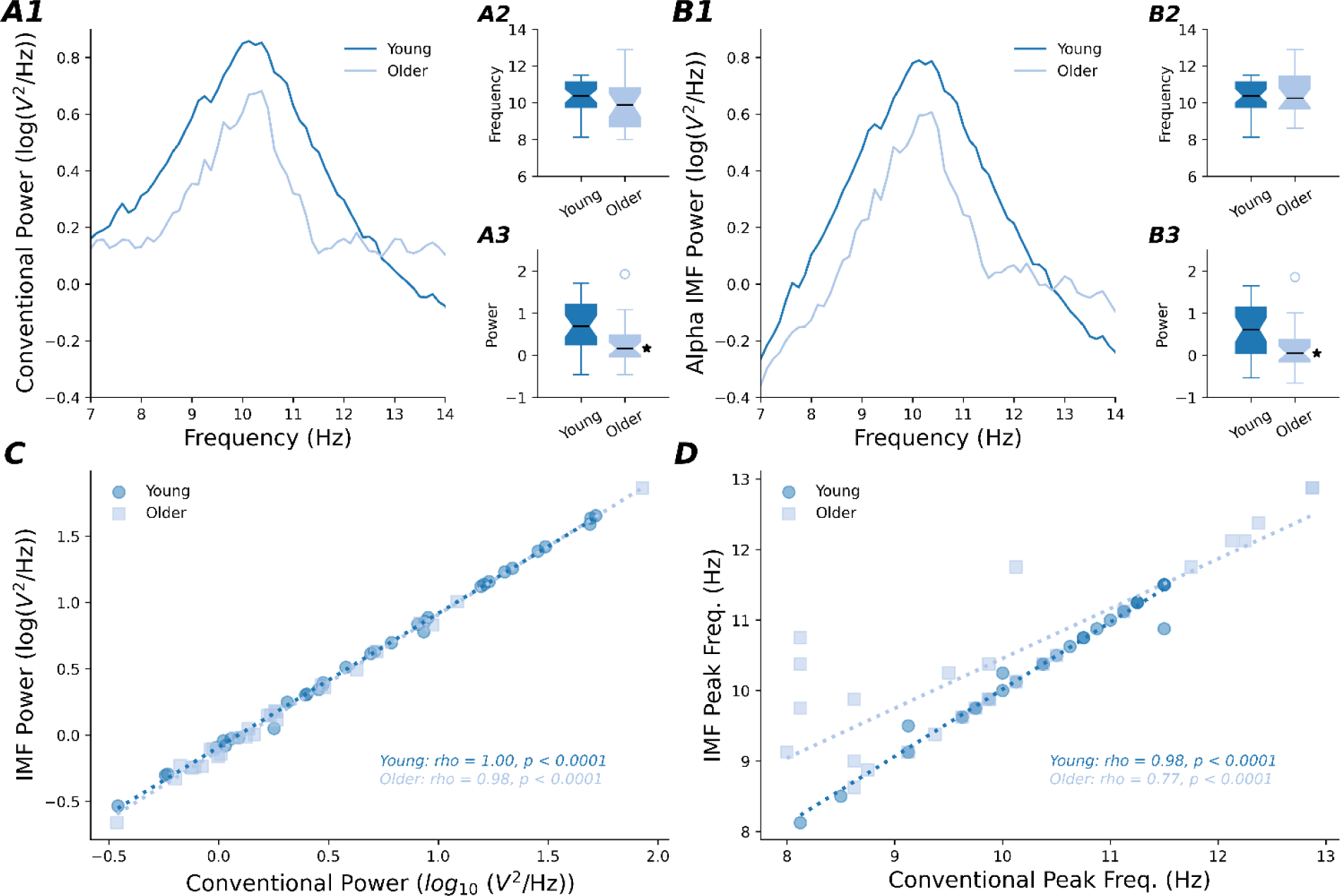
Alpha oscillation characteristics derived from broadband and IMF data. **(A, B)** Alpha band (8-13 Hz) characteristics quantified by application of Welch’s method to broadband data *(A)* and the alpha IMF *(B)*. Power spectral density curves *(A1, B1)*, peak frequency *(A2, B2)*, and power *(A3, B3)* are compared between groups. **(C, D)** Correlations between the estimates of power *(C)* and peak frequency *(D)* derived from each method. *P & 0.05 when compared to the young group.

### Mean within-cycle instantaneous frequency varies between tasks and groups

Table 1 shows IF values for each group, task, and band, averaged over cycle phase. Between-group comparisons of alpha frequency showed higher values in older participants that were consistent (all *pd* > 98.9%) and significant (all 0% in ROPE) for all tasks and time points. For older participants in both GNG and SRT tasks, consistent (all *pd* > 99.4%) and significant (all 0% in ROPE) reductions in frequency were observed during the post-go period, relative to pre-go, post-react, as well as to the rest condition. Furthermore, reductions in frequency relative to the rest condition were consistent and significant for SRT*_pre-go_* (*pd* = 99.3, 0% in ROPE). Also, interaction contrasts suggested that reductions in alpha frequency during the post-go period of the RT task (relative to pre-go. post-react, and rest) and GNG task (relative to post-react only) were significantly and consistently (all *pd* > 99.2%, 0% in ROPE for all comparisons) greater in older compared to young adults. All other within-group comparisons were inconsistent and failed to provide sufficient evidence to accept or reject the null hypothesis (all *pd* < 98.2%, all % in ROPE between 3.3% and 73.2%).

**Table 1.**
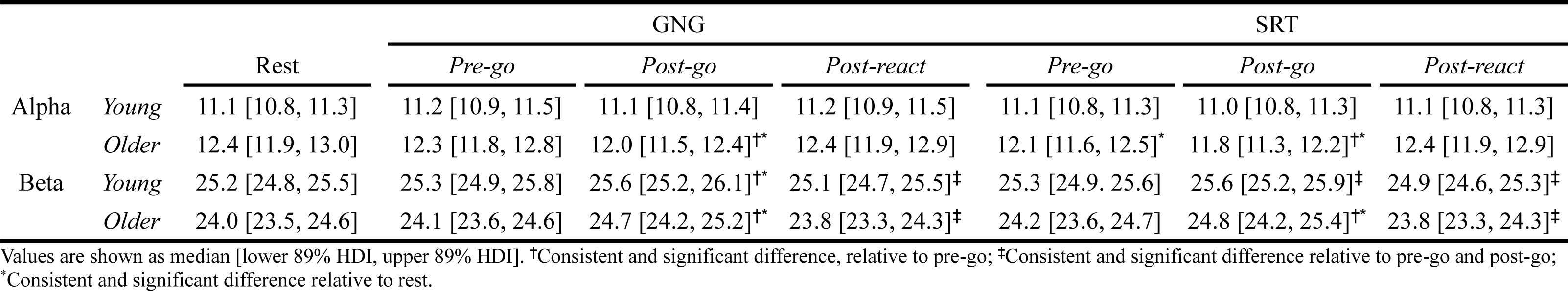
Effects of task on mean within-cycle instantaneous frequency.

Between-group comparisons of beta frequency showed lower values in older participants that were consistent (all *pd* > 96.6%) and significant (all 0% in ROPE) for all tasks and time points. Within-group comparisons for young and older adults separately showed that frequency during both GNG and SRT was increased at the post-go time point, relative to pre-go and post-react, and these differences were consistent (all *pd* > 98.1%) and significant (all 0% in ROPE). For older participants, frequency during post-go was increased relative to rest for both GNG (*pd* = 98.4%, 0% in ROPE) and SRT (*pd* = 98.3%, 0% in ROPE). For young participants, frequency during post-go was significantly increased relative to rest for GNG (*pd* = 98.1%, 0% in ROPE), whereas SRT showed a relatively consistent increase (*pd* = 93.5%) that failed to reach a practical level of significance (4% in ROPE). Furthermore, frequency during the post-react period was reduced relative to pre-go, and this was consistent (*pd* > 99.9%) and significant (all 0% in ROPE) for GNG and SRT in both groups. All other comparisons were inconsistent and failed to provide sufficient evidence to accept or reject the null hypothesis (all *pd* < 84.9%, all % in ROPE between 9.4% and 21.1%).

### Alpha and beta waveforms are non-sinusoidal

Figure 2 shows alpha and beta IF as a function of phase, collapsed over group and task. For alpha cycles, the mean profile suggested that peak frequency of ∼12.4 Hz tended to occur during the zero crossings, but dropped to ∼12.0 Hz during the peak (Fig 2, left panel), consistent with a waveform having a broadened peak. In support of this, examination of mean vectors indicated positive x-axis (mean ± SD = 0.22 ± 0.74) and negative y-axis (mean ± SD = −0.057 ± 1.23) values, both of which were significantly different to zero (x-axis: *t*_204,796_ = 136.5, *P* < 0.0001; y-axis: *t*_204,796_ = −21.3, *P* < 0.0001) (see inset of Fig 2, left panel). For beta cycles, the mean profile suggested that peak frequency of ∼26 Hz occurred during the descending zero crossing but dropped to ∼24.6 Hz just after the trough (Fig 2, right panel), consistent with a waveform having a fast descending edge and broadened trough. Mean vectors for the beta profile were positive (x-axis mean ± SD = 0.29 ± 1.35; y-axis mean ± SD = 0.062 ± 1.62) and significantly different to zero (x-axis: *t*_277,684_ = 113.7, *P* < 0.0001; y-axis: *t*_277,684_ = 20.1, *P* < 0.0001) (see inset of Fig 2, right panel).

**Figure 2.**
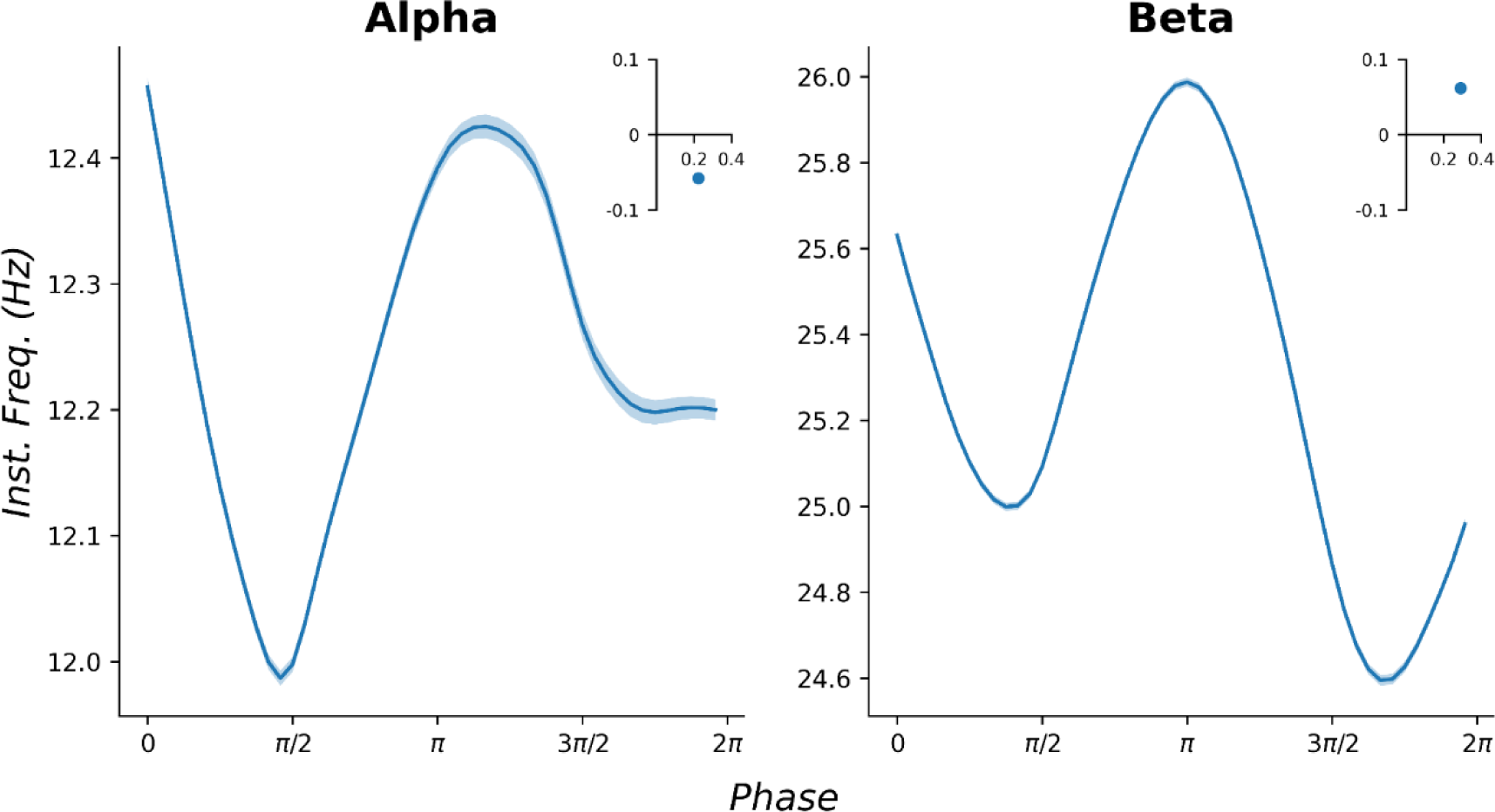
Phase aligned instantaneous frequency. Main panels show mean phase-aligned IF profiles for alpha *(left)* and beta *(right)* modes, whereas insets show mean vectors. Data are collapsed over age group and task; shaded section represents standard error of the mean.

### Waveform motifs are differentially influenced by task and group

Figure 3 shows IF profiles and normalised waveforms for the first 5 components generated by PCA. These explained > 97% of variance in IF for both bands and described increasingly complex elements of waveform shape. For example, while PC1 described opposing variations in the speed of the ascending/descending half waves, and PC2 described opposing variations in the speed of the ascending/descending edges, subsequent PCs instead related to increasingly fractionated facets of cycle phase. Investigation of group, task, and time-point effects on between-cycle variance for each score identified 4 components of interest: alpha PC4 and beta PC1, PC2, and PC3 (Figure 4).

**Figure 3.**
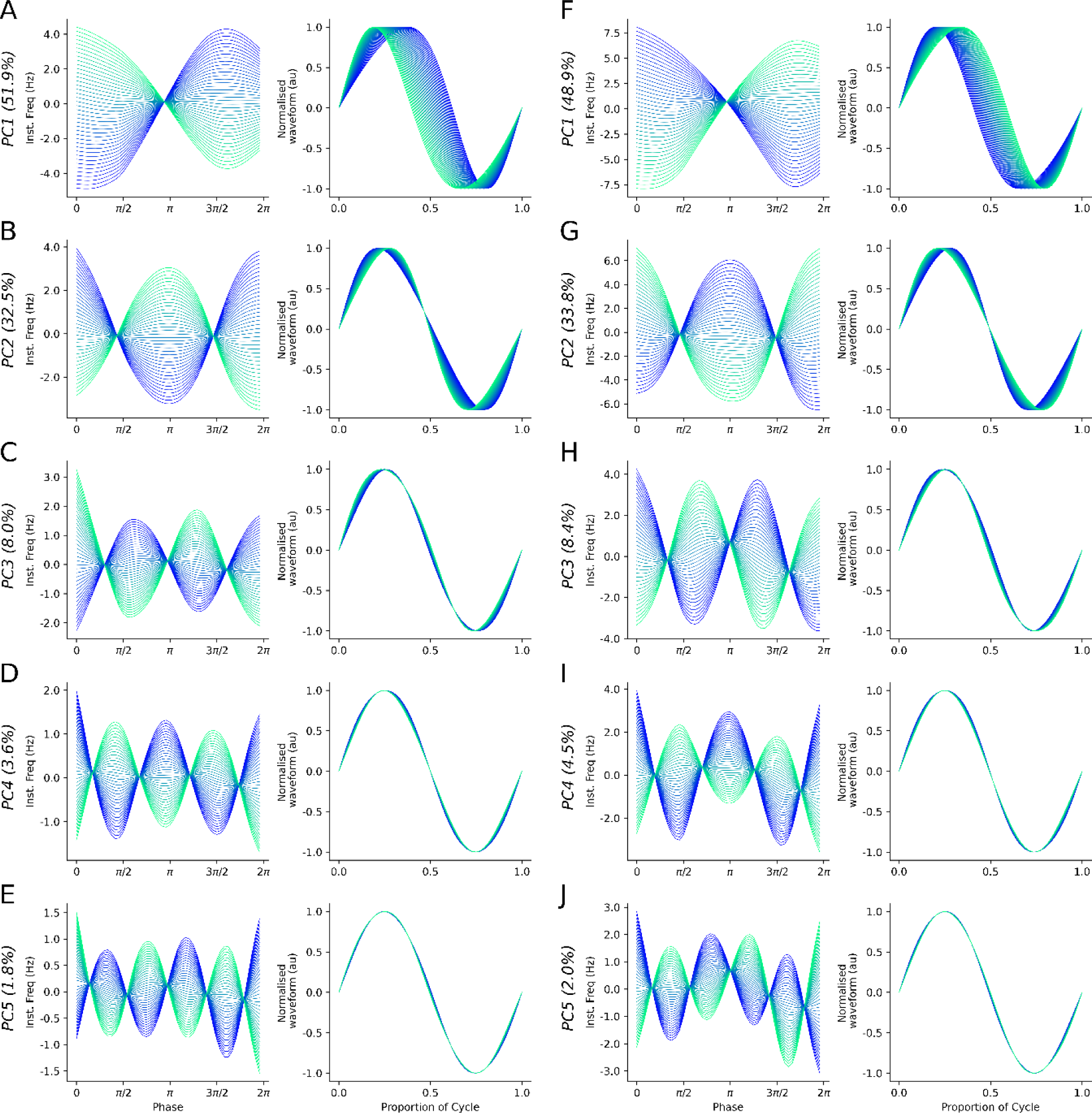
Principal components of waveform shape. ***(A-J, left panel)*** Demeaned IF profiles for the first 5 principal components, plotted from the first percentile (blue lines, negative scores) to the ninety-ninth percentile (green line, positive scores) of scores observed for each component. ***(A-J, right panel)*** Normalised waveforms generated by the IF profiles for each PC score, demonstrating the influence of each PC on waveform shape. Components for the alpha mode are shown in the left column, whereas components for the beta mode are shown in the right column. The bracketed value after the PC label shows the percentage of variance explained by that component.

**Figure 4.**
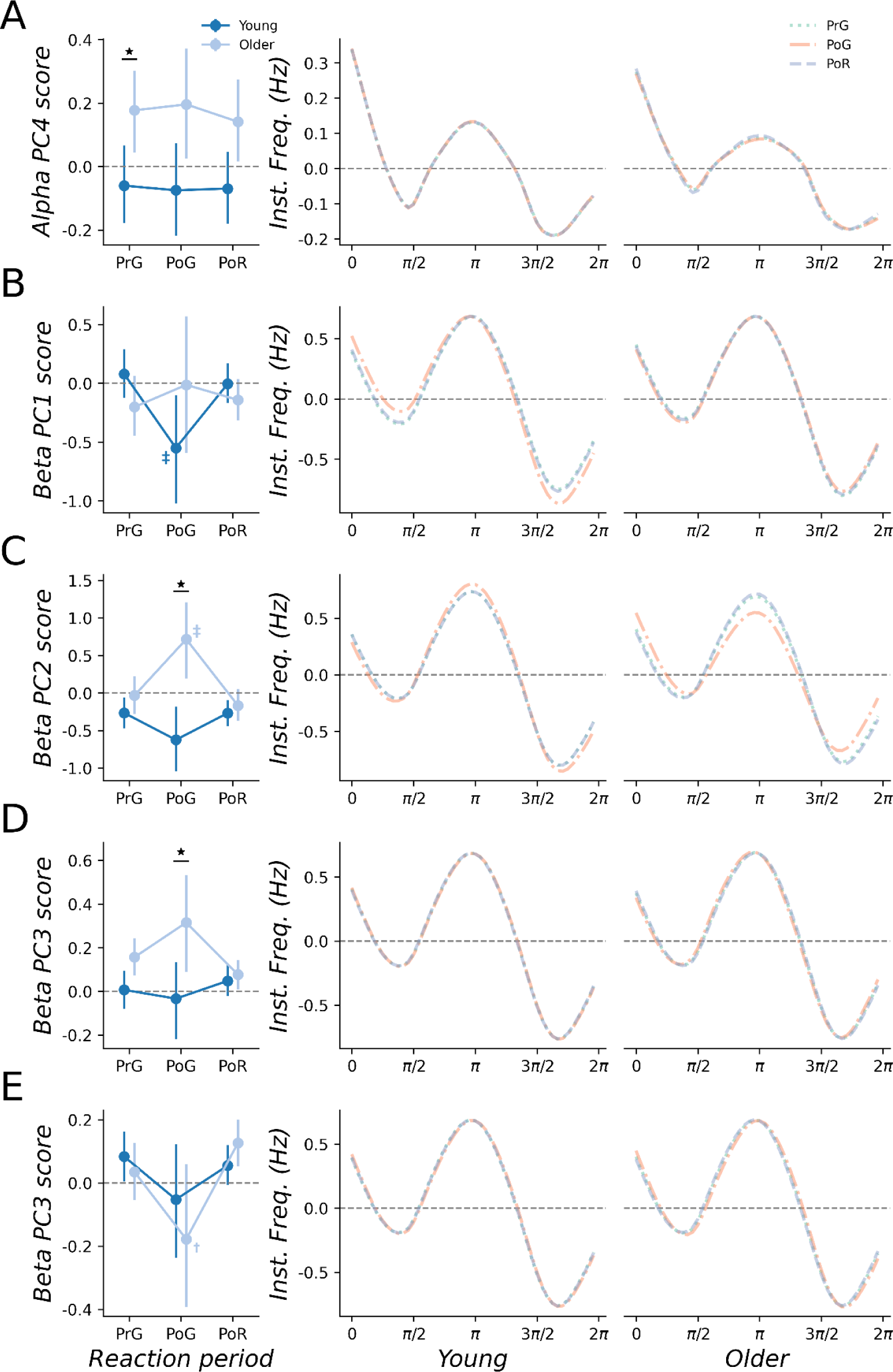
Waveform shape is altered by age and task. **(A-E, left panel)** Median PC scores compared between reaction phases and groups *(young: dark blue; older: light blue)* for the SRT *(A-D)* and GNG *(E)* tasks. Error bars indicate 89% HDI. *Significant and consistent difference between groups; ^†^Significant and consistent difference, relative to post-react. ^‡^Significant and consistent difference relative to both pre-go and post-react periods. **(A-E, right panel)** Projection of median PC scores shown in the right panel. The resulting IF profiles are compared between the pre-go *(dotted green line)*, post-go *(dashed-dotted orange line)* and post-react *(dashed purple line)* time periods in young *(left column)* and older *(right column)* adults.

For alpha PC4 in the SRT task, scores in older adults during the pre-go period were increased relative to the young group (*pd* = 98.1%, 0% in ROPE; Fig 4A, left panel). While increases in older adults were also consistent for the post-go (*pd* = 97.2%) and post-react (*pd* = 97.4%) SRT periods, there was partial overlap with the ROPE (0.4% and 1.5% in ROPE, respectively). All other contrasts were inconsistent and failed to provide sufficient evidence to accept or reject the null hypothesis (all *pd* < 95.34%, all % in ROPE between 17.0% and 97.7%). For beta PC1 in the SRT task, scores in young participants during the post-go period were reduced relative to both pre-go (*pd* = 98.0%, 0% in ROPE) and post-react (*pd* = 97.0%, 0% in ROPE; Fig 4B, left panel). All other contrasts were inconsistent and failed to provide sufficient evidence to accept or reject the null hypothesis (all *pd* < 91.3%, all % in ROPE between 4.87% and 25.4%). For beta PC2 of the SRT task, scores in older participants during the post-go period were increased relative to pre-go (*pd* = 98.6%, 0% in ROPE), post-react (*pd* = 99.5%, 05 in ROPE; Fig 4C, left panel), and rest (*pd* = 99.1%, 0% in ROPE). Furthermore, interaction contrasts showed that increases in score during the post-go period (relative to pre-go, post-react, and rest) were consistently and significantly greater in older adults compared to young adults (all *pd* > 99.6%, 0% in ROPE for all comparisons). In addition, between-group comparisons for the post-go period showed larger scores in older adults that were consistent (*pd* = 99.9%) and significant (0% in ROPE). All other contrasts were inconsistent and failed to provide sufficient evidence to accept or reject the null hypothesis (all *pd* < 93.2%, all % in ROPE between 4.2% and 33.5%). For beta PC3 in the SRT task, scores in older participants during the post-go period were increased relative to the young group (*pd* = 97.6%, 0% in ROPE; Fig 4D, left panel). Between-group comparisons for the pre-go period also showed consistent increases in the older group, but these were not practically significant (4.7% in ROPE). In addition, the older group showed a consistent (*pd* = 95.6%) increase in PC3 scores in post-go relative to post-react, but this was not practically significant (4.0% in ROPE). For PC3 in the GNG task, scores in older adults during the post-go period were reduced relative to post-react (*pd* = 98.1%, 0% in ROPE; Fig 4E, left panel) and rest (*pd* = 97.5%, 0% in ROPE). All other contrasts were inconsistent and failed to provide sufficient evidence to accept or reject the null hypothesis (all *pd* < 97.6%, all % in ROPE between 4.0% and 61.0%).

### Correlation analysis

To investigate the relationship between waveform shape and motor function, RT was correlated against PC scores within each time point (quantified via median values and the coefficient of variation) using robust Bayesian correlations. This approach revealed correlation coefficients ranged from −0.095 to 0.136 across all comparisons, with a median value of −0.002. Consequently, RT values were unrelated to median scores, or the variance in scores, within each time point.

## Discussion

Oscillatory cycles within the brains electrical activity are typically described by their peak frequency or amplitude. However, a developing literature suggests that examination of waveform shape provides additional insight into the physiological and functional relevance of oscillatory activity. Within the current study, we aimed to assess if waveform shape provides novel information about how the ageing process alters the brain, particularly with respect to motor function. To achieve this, we applied a recently developed approach, where within-cycle fluctuations in IF index complex waveforms that are subsequently decomposed into motifs explaining different dimensions of shape. Using this approach, we identified task- and age-dependent changes in the shape of oscillatory cycles occurring within the alpha and beta range.

### Alpha and beta cycles exist across a range of waveform shapes

Quantification of oscillatory activity has been a key element of EEG and magnetoencephalography (MEG) research since the inception of these techniques, with their examination now supported by dedicated and extensive fields of research. These have established important contributions of oscillations to numerous functions within the brain, including communication between areas (Fries 2005, Fries 2015), integration of information across different spatiotemporal scales (Canolty and Knight 2010, Bonnefond *et al*. 2017) and gating of information (Jensen and Mazaheri 2010). While the methodology available for the investigation of oscillatory activity is extensive, some variant of spectral analysis is generally applied. This often involves Fourier or wavelet convolution to decompose the broadband signal into narrowband frequency components, with changes in the power and peak frequency within canonical bands taken to reflect altered oscillatory activity (Donoghue *et al*. 2022). However, these convolution techniques assume a sinusoidal waveform. In contrast, both recent and historical literature recognises that oscillatory activity is frequently non-sinusoidal (for review, see; Cole and Voytek 2017). For example, the sensorimotor mu rhythm demonstrates an arch or wicket shape (Gastaut 1952, Kuhlman 1978), whereas the sensorimotor beta rhythm has been shown to have a sawtooth waveform (Cole et al. 2017, Jackson et al. 2019), as too have theta (Belluscio *et al*. 2012, Ghosh *et al*. 2020, Quinn et al. 2021b) and gamma (Krishnakumaran *et al*. 2022) oscillations.

Within the current study, we examined individual cycles of oscillatory activity with frequencies peaking in the alpha and beta range. For both, investigation of mean vectors suggested non-zero values that indicate a departure from a sinusoidal shape (Quinn et al. 2021b). Consequently, our results further support the non-sinusoidal nature of these brain rhythms. However, examination of IF profiles for both bands suggest relatively increased speed of edges and decreased speed of extrema, which contrasts with previous descriptions of feature asymmetries (i.e., fast ascending edge/peak-slow descending edge/trough)(Cole et al. 2017, Cole and Voytek 2017). Despite this, the magnitude of these shifts was not consistent across cycle features; for example, the decrease in alpha IF was greater at the peak than the trough, whereas the decrease in beta IF was greater at the trough than the peak (Fig 2). Furthermore, mean vectors showed substantial variability around the average (see Results), including examples across all parts of the complex plane describing edge (e.g., x-axes in insets of Fig 2) and extrema (e.g., y-axes in insets of Fig 2) asymmetries. Taken together, IF profiles and mean vectors suggest that alpha and beta cycles exist across a wide range of shapes. This is consistent with recent findings for oscillations in theta (Quinn et al. 2021b) and beta (Szul et al. 2022, Rayson et al. 2023) bands.

To characterise the different facets of waveform shape, we implemented a recently developed approach involving identification of waveform motifs via PCA-based decomposition across individual cycles (Quinn et al. 2021b, Szul et al. 2022, Rayson et al. 2023). The first 5 PCs from this analysis explained more than 97% of variance in waveform shape for both alpha and beta modes, with the majority of this partitioned within PC1 (alpha = 51.9%, beta = 48.9%) and PC2 (alpha = 32.5%, beta = 33.8%). For both modes, these described inversely related fluctuations in the speed of the first and second half wave (PC1), and ascending and descending edges (PC2) (Fig 3). Consequently, the first two PCs can be considered to characterise the dimensions of waveform shape indexed by control point-based measures of peak-trough and rise-decay asymmetry, respectively (Cole et al. 2017, Cole and Voytek 2019). Importantly, these components recover descriptions of waveform shape that are consistent with previous studies but were not apparent in the composite IF profile (see preceding paragraph). Subsequent PCs explained substantially less variance; despite this, they nonetheless provide additional insight into increasingly subtle fluctuations in waveform shape, some of which were uniquely influenced by age and/or task (see below), demonstrating the potential value of the information they contain.

### Age and task uniquely influence specific facets of waveform shape

Scores associated with each PC described the extent to which different waveform motifs contributed to each cycle. Comparing scores between age groups and task conditions therefore allowed us to examine if/how each dimension of waveform shape is sensitive to different motor paradigms, and if the nature of this relationship is altered by advancing age. These comparisons identified several PCs that were significantly modulated (Fig 4), revealing differences between groups that were dependent on reaction phase (e.g., beta PC2), in addition to those that were not (e.g., alpha PC4). Consequently, our results suggest that examining waveform shape in this way provides novel insight into the neurophysiological effects of ageing within the motor system. A caveat to this conclusion is that PC scores were unrelated to motor performance assessed via reaction time, which limits the functional implications we can infer. However, several factors could have contributed to this lack of correlation, and we do not believe it demonstrates functional irrelevance per se. For example, using a similar PCA-based approach, two recent studies showed that task-dependent changes in beta bursts were limited to specific centiles of PC scores, with greater modulation apparent in scores further from the median (Szul et al. 2022, Rayson et al. 2023). Consequently, examination of different points within the spectrum of shape described by each PC (e.g., Fig 3) may have provided additional information with respect to functional relevance. We decided against similar analyses within the current study, mostly because the narrow post-go time window (within which the largest effects were apparent) only allowed identification of a relatively small number of cycles which would not be amenable to further subdivision. A related point is that both pre-go and post-react periods were relatively prolonged, which may have obscured changes in shape related to task performance. Our decision to examine wide time windows stemmed from the exploratory nature of this project and our desire to maximise the number of oscillatory cycles available for analysis. Future work will therefore need to examine specific time windows that can be expected to relate more directly to task performance.

Alternatively, it may be that the neurophysiological processes contributing to the examined dimensions of waveform shape are unrelated to the elements of performance indexed by reaction time, which represents a relatively gross measure of motor function. Our decision to use open-source data for this project meant that the way in which performance was assessed was largely predetermined. However, this point clearly demonstrates the need for further examination in future studies using alternative motor tasks. It will be of particular benefit to implement tasks that include extended reaction periods where the temporal evolution of waveform shape can be examined in greater detail. This is supported by our supplementary plots of PC scores over time; while the reduced sample density within the post-go period means these must be interpreted with caution, they nonetheless demonstrate substantial fluctuations not captured by the main analysis (see supplementary figures S2-S5).

While the neurophysiological processes driving waveform shape are not well understood, the developing literature has identified several contributing mechanisms. For example, seminal work used multimodal evidence (local field potentials, MEG, modelling) in animal and human data to show that the shape of somatosensory beta bursts are driven by synchronous thalamic inputs to proximal and distal dendrites of pyramidal neurons (Sherman *et al*. 2016), and recent work with MEG supports a similar mechanism in human motor cortex (Bonaiuto *et al*. 2021). Furthermore, Cole and Voytek (2018) used hippocampal recordings in rodents to show that non-sinusoidal features of theta cycles relate to synchronisation, activation sequence, and firing rate in pyramidal neurons and interneurons; recent work from Garcia-Rosales *et al*. (2023) also demonstrated the importance of synchronisation within local spiking activity for driving the shape of delta cycles within the fronto-auditory circuit of bats. Finally, Marshall *et al*. (2022) recently showed that application of transcranial direct current stimulation changes one half-wave of gamma cycles (recorded from human visual cortex with MEG), and used biophysical modelling to show that altered excitability of layer 5 pyramidal cells could explain their results. Taken together, this literature demonstrates that waveform shape reflects the structure and dynamics of local cortical networks, and their modulation by interconnected areas. The details of this largely remain to be determined, and likely reflect varied processes across different brain areas (e.g., Garcia-Rosales et al. 2023), making it difficult to speculate about their relation to the specific changes in waveform shape we report here. Nonetheless, it seems reasonable to suggest that our results may reflect task- and age-dependent changes in cortical network activity/connectivity. Further exploration of the mechanisms underpinning variations in waveform shape, possibly via pharmaceutical intervention or the application of non-invasive brain stimulation, will be critical for understanding the effects reported here.

### Instantaneous frequency is altered in older adults

Age-related comparisons of IF suggested that older adults showed an increase in cycle frequency for the alpha mode but decrease in cycle frequency for the beta mode. These outcomes contrast with previously reported effects of older age on the peak frequency of alpha and beta oscillations (see *Introduction*), derived using Fourier-based methods. Given that, within the current study, estimates of peak frequency using Welch’s method were highly comparable for both the raw data and the alpha IMF (Fig 1), this outcome is unlikely to reflect an artifact of EMD. It is also important to note that phase-aligned IF data were demeaned prior to PCA, meaning that our analysis of waveform motifs was not influenced by between-group differences in IF. That being said, Fourier-based estimates of peak frequency and Hilbert-Huang based estimates of IF cannot be considered to provide the same information about oscillatory activity (Huang et al. 1996, Huang et al. 1998, Huang et al. 2009). In particular, Fourier based methods assume a linear and stationary signal (in contrast to the non-stationary nature of EEG), whereas estimates of IF are sensitive to the intra- and inter-wave fluctuations which are present in EEG data (Lo et al. 2009). Consequently, previously reported effects of age on oscillatory frequency (as calculated using Fourier-based techniques) have a temporal resolution that is limited by the length of the time window over which the Fourier transform was applied. In contrast, the IF data reported here were estimated directly from the narrowband IMFs via the differentiation of phase, and therefore resolve fluctuations in frequency on a much higher temporal scale (i.e., the level of individual oscillatory cycles). In addition to differences in temporal resolution, these techniques are also differentiated by the way in which they isolate oscillatory activity: conventional approaches using Fourier methods quantify activity within *a priori* bands, whereas analyses of IF can include a much broader range of frequency values. Taken together, the previous and current examinations of age-related changes in oscillation frequency likely interrogate different characteristics of the underlying generators, possibly relating to different timescales of variability.

In conclusion, the current study examined how the shape of oscillations within the alpha and beta bands varied in young and older adults as they performed SRT and GNG reaction time tasks. To achieve this, we implemented an analysis pipeline that characterised waveform shape based on within-cycle fluctuations in phase-aligned IF, which were subsequently decomposed into waveform motifs using PCA. Comparisons of motif scores identified several dimensions of shape that were influenced by age and/or task, possibly providing unique insight to cortical activity and connectivity associated with motor function in young and older adults. Future work that further examines how these fluctuations in shape relate to performance, in addition to the underlying mechanisms, will be important for identifying how this information can be used to understand age-related changes in the brains motor networks, and to inform new methods for influencing motor function in the elderly.

## Acknowledgements

The authors would like to thank Dr Andrew Quinn for useful advice on implementation of the waveform analysis pipeline. GMO was supported by a discovery early career research award from the Australian Research Council (DE230100022).

## Data availability

Pre-processed EEG data and code used within analyses of the current study is available from https://osf.io/ctq98/.

