## supplementary figures for "Healthy ageing influences how the shape of alpha and beta oscillations change during reaction time tasks"

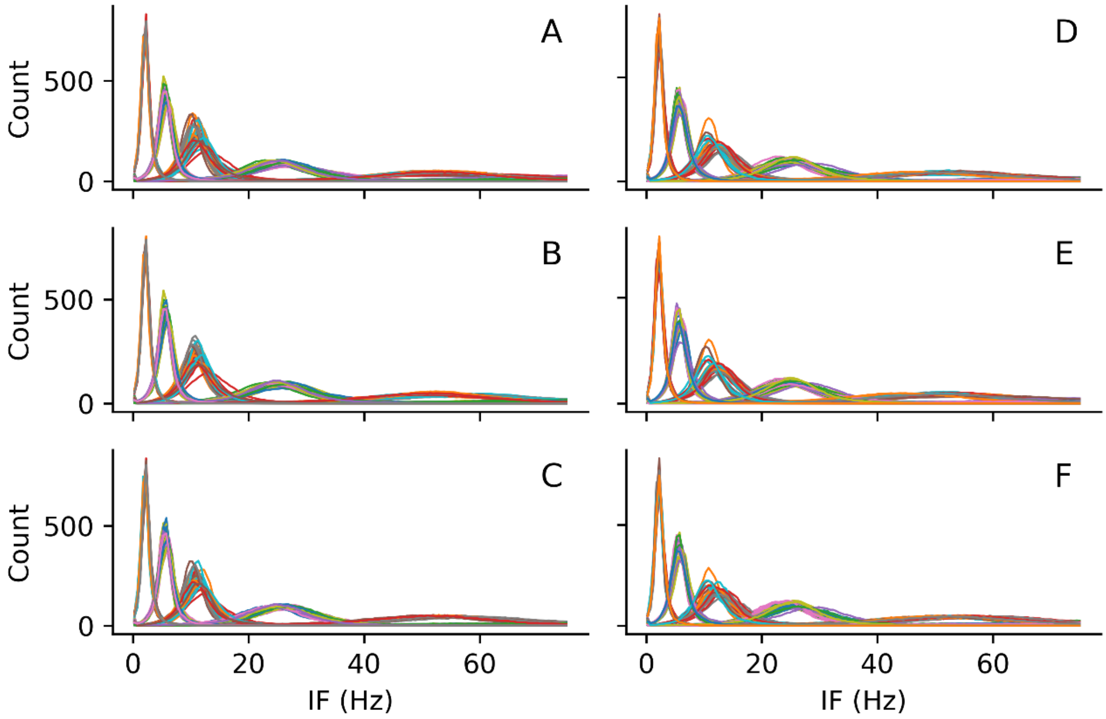


**Figure S1.** Comparison of IMFs generated within individual young *(left column)* and older *(right column)* participants during rest ***(A, D)***, SRT ***(B, E)*** and GNG ***(C, F)*** tasks.

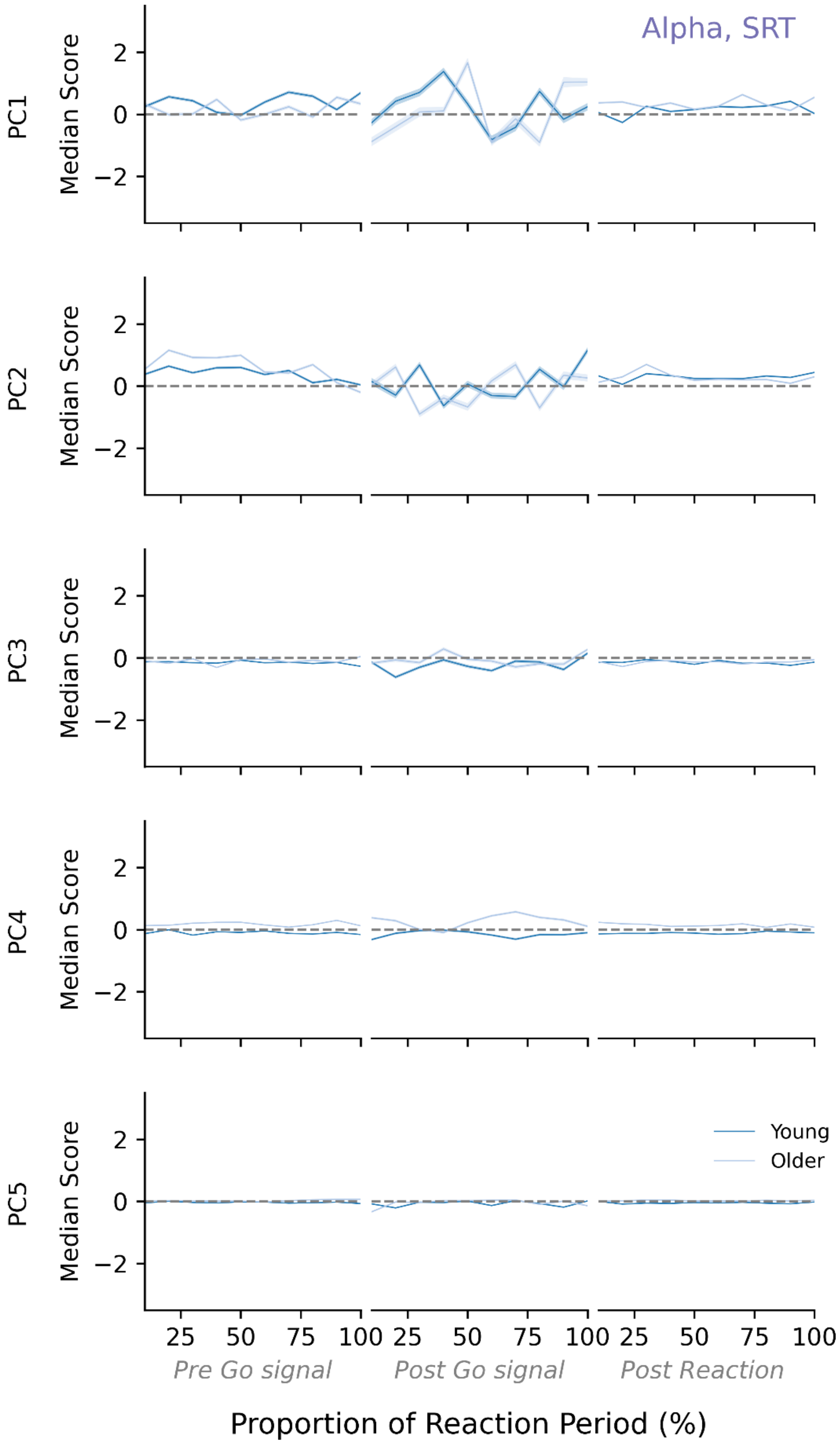


**Figure S2.** Temporal fluctuations in the first 5 PCs of the alpha waveform during the pre-go *(left column)*, post-go *(middle column)* and post-react *(right column)* time points of the SRT task. Values are compared between young *(dark blue)* and older *(light blue)* adults, timed relative to each reaction phase. Shaded section indicates standard error of the mean.

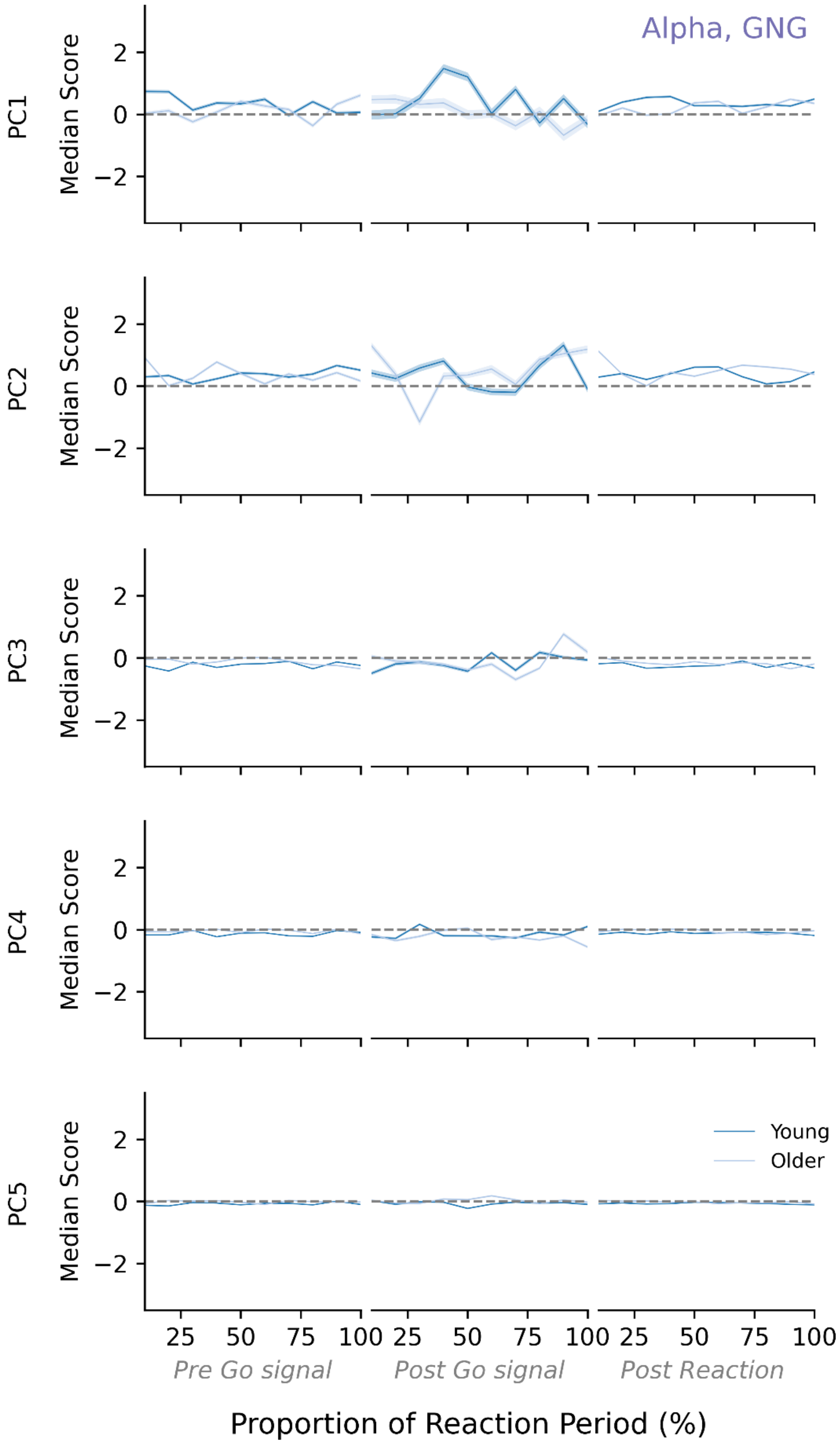


**Figure S3.** Temporal fluctuations in the first 5 PCs of the alpha waveform during the pre-go *(left column)*, post-go *(middle column)* and post-react *(right column)* time points of the GNG task. Values are compared between young *(dark blue)* and older *(light blue)* adults, timed relative to each reaction phase. Shaded section indicates standard error of the mean.

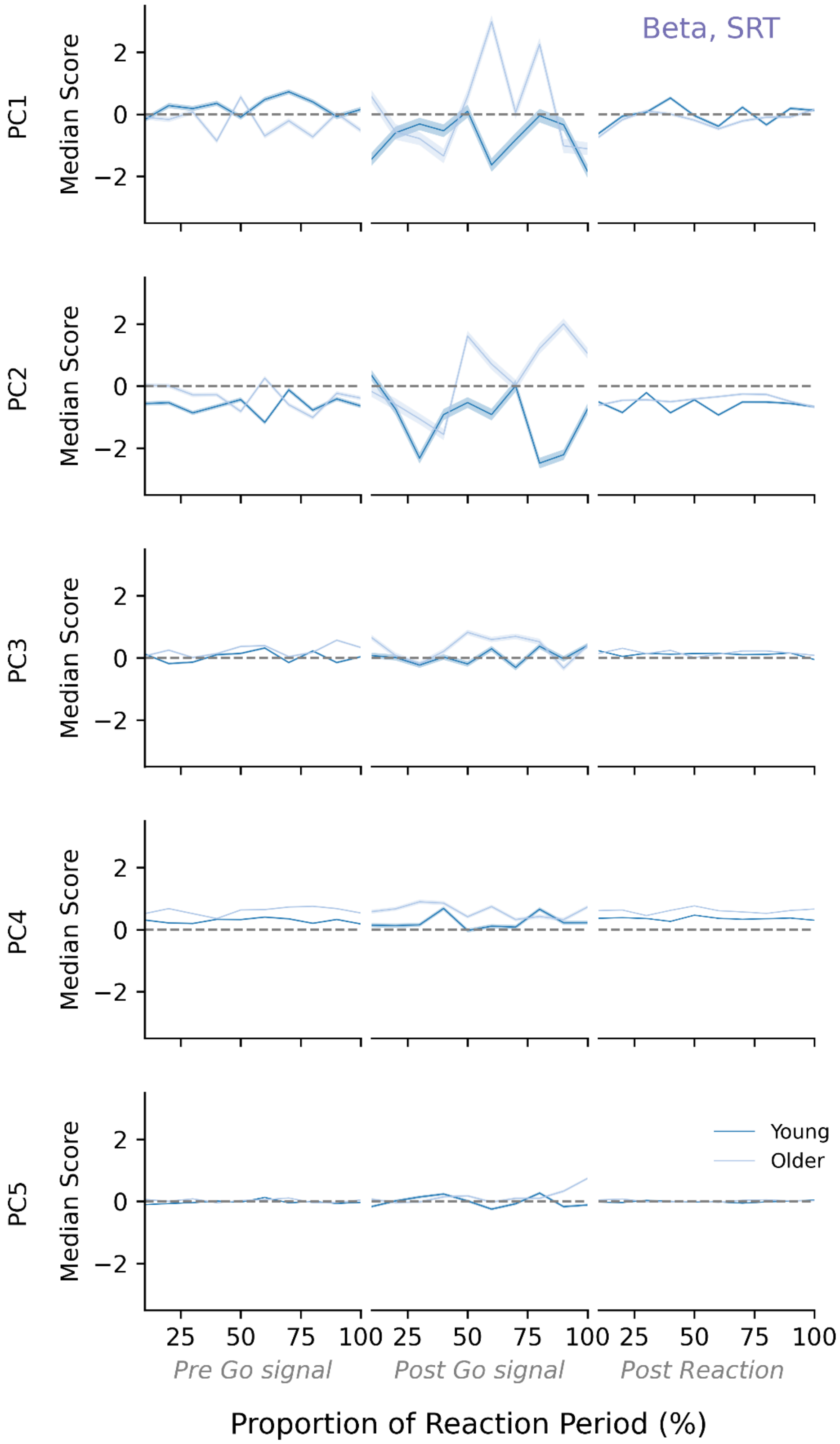


**Figure S4.** Temporal fluctuations in the first 5 PCs of the beta waveform during the pre-go *(left column)*, post-go *(middle column)* and post-react *(right column)* time points of the SRT task. Values are compared between young *(dark blue)* and older *(light blue)* adults, timed relative to each reaction phase. Shaded section indicates standard error of the mean.

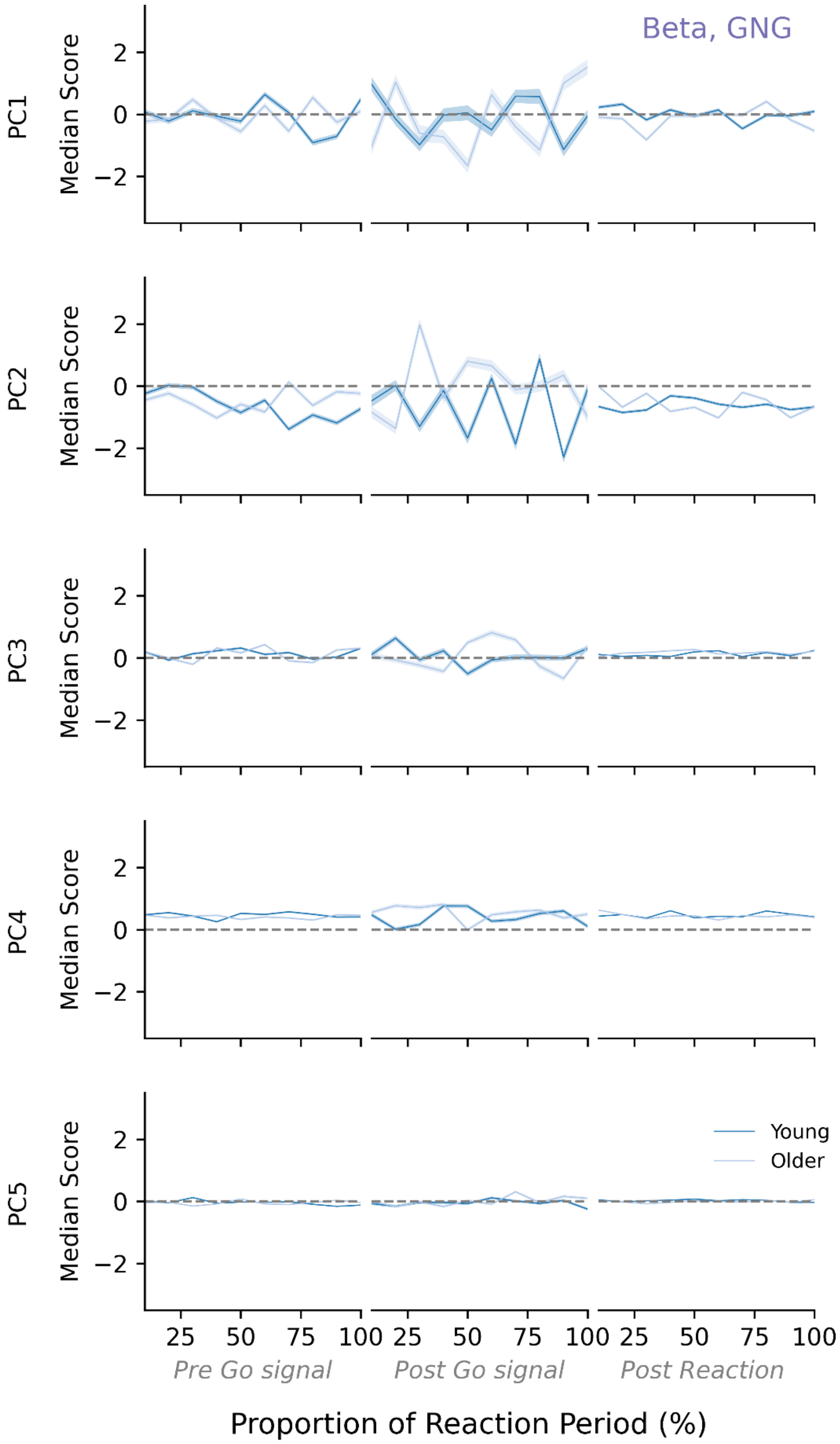


**Figure S5.** Temporal fluctuations in the first 5 PCs of the beta waveform during the pre-go *(left column)*, post-go *(middle column)* and post-react *(right column)* time points of the GNG task. Values are compared between young *(dark blue)* and older *(light blue)* adults, timed relative to each reaction phase. Shaded section indicates standard error of the mean.
